# Improving HEK293-based AAV-production using GSMMs, and a multi-omics approach

**DOI:** 10.1101/2024.10.10.617556

**Authors:** L. Zehetner, D. Széliová, B. Kraus, J. A. Hernandez Bort, J. Zanghellini

## Abstract

HEK293 cells are a versatile cell line extensively used in the production of recombinant proteins and viral vectors, notably Adeno-associated virus (AAV) [12]. Despite their high transfection efficiency and adaptability to various culture conditions, challenges remain in achieving sufficient yields of active viral particles. This study presents a comprehensive multi-omics analysis of two HEK293 strains under good manufacturing practice conditions, focusing on the metabolic and cellular responses during AAV production. The investigation included lipidomic, exometabolomic, and transcriptomic profiling across different conditions and time points. Genome-scale metabolic models (GSMMs) were reconstructed for these strains to elucidate metabolic shifts and identify potential bottlenecks in AAV production. Notably, the study revealed significant differences between a High-producing (HP) and a Low-producing (LP) HEK293 strains, highlighting pseudohypoxia in the LP strain. Key findings include the identification of hypoxia-inducible factor 1-alpha (HIF1alpha) as a critical regulator in the LP strain, linking pseudohypoxia to poor AAV productivity. Inhibition of HIF1alpha resulted in immediate cessation of cell growth and a 2-fold increase in viral capsid production, albeit with a decreased number of viral genomes, impacting the full-to-empty particle ratio. This suggests that while HIF1alpha inhibition enhances capsid assembly, it simultaneously hampers nucleotide synthesis via the pentose phosphate pathway (PPP), necessary for genome packaging.

## Introduction

Human embryonic kidney 293 (HEK293) cells represent a highly adaptable cell line originating from human embryonic kidney cells cultivated in tissue culture. Initially developed in 1977 [29], HEK293 cells have since become pivotal in the generation of recombinant proteins [69, 23, 46, 1] and viral vectors [12, 65, 14, 81], owing to their transfection efficiency and robust proliferation characteristics. Various databases, such as Cellosaurus [4], document 564 distinct HEK293 strains, each optimized for specific applications. These cells are extensively used in the biopharmaceutical sector for the production of recombinant proteins that are not effectively expressed in Chinese Hamster Ovary (CHO) cells [45]. A significant advantage of HEK293 cells is their adaptability to diverse culture conditions; they are capable of growing in both suspension and adherent cultures [46, 42] and are suitable for serum-free media, which facilitates scalable and reproducible production processes.

HEK293 cells are extensively utilized for the production of various viral vectors essential for cell and gene therapy [9, 56, 39], as well as vaccine development [38, 15]. These vectors include lentiviruses, adenoviruses, and AAV [12]. AAV, in particular, have gained significant prominence for *in vivo* gene therapy, with six drugs receiving approval in recent years [14]. Despite their considerable therapeutic potential, the production of sufficient quantities of active viral particles remains challenging due to high associated costs, and the fact that only 10% of the produced AAV contain the gene of interest in full length [49, 48]. To mitigate this challenge, numerous studies have aimed at enhancing AAV titers in HEK293 cells [9, 19]. Beyond optimizing process parameters, several investigations have examined the cellular behavior of HEK293 cells using single-omics datasets during growth [46, 23, 47] or production [45, 65, 23, 58, 64, 75, 56], as summarized in [1].

Focusing on AAV particles, the host cell behavior during AAV production has been extensively studied using transcriptomes [75, 56, 39], proteomes [65, 64], or both [43] (Tab 1). Although fermentation conditions, strains, and AAV serotypes varied across studies, two pathways were consistently identified as enriched in the production state: ER stress and the unfolded protein response (UPR) [39, 56, 75, 43, 65], and immune response [64, 39, 75, 43]. The accumulation of unfolded or misfolded proteins in the endoplasmic reticulum (ER) induces ER stress, triggering the UPR to restore ER homeostasis. Key proteins involved in UPR activation include GRP78, ATF6, PERK, and IRE1. PERK kinase phosphorylates *eIF*2*α*, resulting in the inhibition of CAP-dependent protein translation [44]. ER stress and UPR are common challenges in bioprocesses for recombinant protein production [13, 57], making the upregulation of the UPR in HEK293 strains during AAV production unsurprising. Additionally, transcriptomic analyses have identified the JAK-STAT signaling pathway as central in HEK293 strains during AAV production [75, 39]. The JAK/STAT pathway is a key mediator of the innate immune response against viral infections and controls the expression of numerous genes involved in antiviral defense, inflammation, cell growth, and differentiation [34, 53, 27]. Subsequent experiments demonstrated that inhibiting STAT1 with Ruxolitinib led to a two-fold titer increase in shake flasks and a 50% improvement in a 3L bioreactor [39]. However, in certain HEK293 strains [39], the JAK-STAT pathway was upregulated even before triple transfection, indicating a pre-existing stressed state, while in other strains, the immune response was either triggered by AAV production [75] or downregulated by ethanol addition [64]. The exact cause of the immune response – whether due to HEK293 strain adaptation or AAV particle production – remains unclear.

**Table 1.** Overview of recently performed -omics based studies of HEK293 strains during AAV production.

| HEK293 strain | Culturing condition | Data | Resource |
| --- | --- | --- | --- |
| HEK293 (ATCC: CRL-1573) | AMBIC media, shake flask | Transcriptome | [75] |
| HEK293 (Mass Biologics, Inc.) | BalanCD media, shake flask | Transcriptome | [75] |
| HEK293 (unknown origin) | FreeStyle F17 media, 50L bioreactors | Transcriptome | [17] |
| HEK293 (FreeStyle 293F) | BalanCD media, shake flask | Proteome | [65] |
| HEK293 (Cell Biolabs, Inc.) | DMEM (+10% FBS), T-flask | Transcriptome, Proteome | [43] |
| HEK293 (Expi293) | HE400AZ media, Erlenmeyer flask | Transcriptome | [64] |
| HEK293 (ATCC: CRL-1573), LP-strain | FreeStyle F17 media, 250mL bioreactors | Transcriptome | [56] |
| HEK293, HP-strain | FreeStyle F17 media, 250mL bioreactors | Transcriptome | [56] |

A potential reason for the immune response of certain strains prior to transfection might be differences between suspension and adherent culturing of HEK293 strains. Recent investigation shed light on the metabolic differences after the adaption of an adherent HEK293 strain to suspension [36]. The study found increased glucose uptake, lactate secretion, and succinate production in HEK293 cells adapted to suspension culture, suggesting that these cells experience pseudohypoxia [31]. In CHO cells, reducing pseudohypoxia by inhibiting pyruvate dehydrogenase kinase with dichloroacetate has been shown to improve culture performance and antibody production [11].

Aside from these efforts, modeling AAV production in HEK293 cells on a genome scale has been largely overlooked to date. GSMMs are mathematical representations of cellular metabolism, with the first GSMM developed over two decades ago [25]. This advancement has led to the creation of thousands of GSMMs across all domains of life [30]. In the context of bioprocesses, GSMMs have provided profound insights at the cellular level and have been successfully applied to both microbes [21, 71, 79, 3] and mammalian cells [35, 16, 55] to enhance growth and production. Currently, no GSMM exists for HEK293 cells during the production of Recombinant protein (RP) or viral particles. Previous models for HEK293 cells have typically been small-scale, focusing on selected pathways [32], or the production pathway of AAV via triple transfection [52], as recently summarized [81]. The most comprehensive HEK293 GSMM for bioprocesses to date encompasses 357 reactions and was manually curated for RP production [58], using transcriptomic and exometabolomic data [23], based on the Recon2 model [66]. In the interim, several updated versions of human metabolism have been published [59, 10], which can be utilized with reconstruction tools to automatically generate cell-specific GSMMs [37, 7, 2, 74, 62].

In this study, we conducted a multi-omics analysis of two HEK293 strains, which were partially examined in previous research [56, 39] for AAV production. Using the latest reconstruction of human metabolism [59], we reconstructed 20 condition-specific GSMMs from the two HEK293 strains, each in two conditions (mock and transfected) across five time points with 4 biological replicates. From GSMMs, we observed that over 99.8% of the resources are used for growth instead of AAV production in both strains. To reallocate the resources, we aimed to inhibit growth and therefore increase AAV production in the LP strain. With the multi-omics analysis – encompassing lipidomes, transcriptomes, and fluxomes – we identified HIF-1*α* as a potential target in the LP strain, suggesting that the LP strain is affected by pseudohypoxia due to recent adaption from adherent to suspension culturing. Inhibiting HIF-1*α* halted cellular growth and improved the specific productivity of viral capsids by over two-fold. However, the number of viral genomes decreased following HIF-1*α* inhibition, leading to a less favorable full-empty ratio. Despite this, our study provides a comprehensive multi-omics analysis and represents a significant advancement in data-driven process design.

## Materials and Methods

### Bioprocess and Sampling

The bioprocess from cell culture to fermentation was performed as described recently by [56]. Briefly, two distinct HEK293 suspension cell line strain stocks, stored at -130°C, were cultured at +37°C in chemically defined, serum-free medium (FreeStyle F17, ThermoFisher, NY, USA) supplemented with 8 mM L-Glutamine and 1.0 g L^−1^ Lutrol. The cultivation was carried out in a HERA Cell 150 (Thermo Fisher Scientific) incubator with 5% CO_2_. To enhance statistical analysis, each cell line was cultivated in four biological replicates using Single Use Spinner flasks (Corning, Germany), with volumes ranging from 61 mL to 1.600 mL during the cell expansion phase. After 12 days, the cells were transferred to an ambr250 Modular fermentation system (Sartorius, Germany) under controlled conditions (pH, CO_2_, O_2_) for transient AAV 8 particle production. Prior to transfection, cells were diluted to a uniform density of 4.0.10^6^ cells per mL to facilitate comparison. Each biological replicate was divided into two fermentation vessels: one for plasmid transfection and the other for mock transfection (without plasmids). A triple-plasmid transfection was performed using Polyethylenimine (Merck KGaA, Darmstadt, Germany), containing Adenovirus 5 Helper genes, Rep2Cap8, and a transgene sequence, in accordance with the supplier‘s instructions. Post-transfection, cells were cultured for 72 hours at +37°C and sampled for analysis at five different time points: before (0 Hours post transfection (HPT)) and after transfection (4, 24, 48, and 72 HPT), resulting in a total of 80 samples (see Fig 1). Additional samples were taken at 21, 27, 45, and 51 HPT for quantification of lactate, ammonia (BioHT), proline, and glycine.

**Figure 1:**
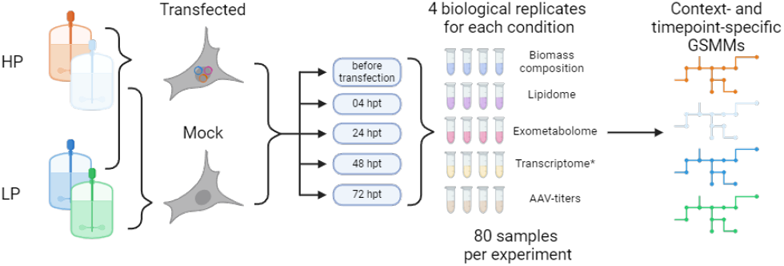
Graphical overview of the methods section. All samples were measured as described in the methods. Transcriptomic data were obtained from [56]

### Experimental Analyses

#### Biomass and cell count

As described in [67], 1 mL of cell suspension was centrifuged at 200 g for 10 minutes at +4°C. The pellet was then washed with 1 mL of pre-cooled phosphate-buffered saline (PBS, 1M, +4°C). To protect the cells during subsequent steps, Pluronic™ F-68 Non-ionic Surfactant (Gibco™, MA, USA) was added to the suspension at a final concentration of 2% (v/v). Glass beakers and silica beads were prepared by drying them overnight at +90°C. They were subsequently cooled in a vacuum desiccator for 20 minutes before being weighed to ensure accuracy. Each sample, resuspended in a 1M PBS solution, was transferred to the pre-weighed beakers. The samples were then dried at +90°C until the biomass remained constant, ensuring complete removal of moisture. Before each weighing, the beakers were cooled for 20 minutes in the vacuum desiccator to prevent heat-related measurement inaccuracies. Cell counts were performed using a NucleoCounter202 (Allerod, Denmark), adhering strictly to the manufacturer’s protocol.

#### Cellular glucose content

In alignment with the methodology described by [67], we posited glucose as the predominant carbohydrate component within the cells and thus limited our carbohydrate quantification to glucose. For this purpose, we employed the Total Carbohydrate Quantification Assay Kit (ab155891, Abcam, Cambridge, UK). Beginning with cell pellets containing 3 − 10·10^6^ cells, we adjusted the volume of lysis buffer proportionately. Sample preparation was carried out following the manufacturer’s instructions. Each sample was analyzed in technical triplicates to enhance the reliability of our measurements.

#### Cellular DNA and RNA content

DNA and RNA content was extracted from the same sample using the AllPrep DNA/RNA Mini Kit (Qiagen, Venlo, NE). As suggested by the manufacturer, *β*-mercaptoethanol (Thermo Fisher Scientific Inc., Waltham, MA, US) was added to inhibit RNases. Samples were quantified on a Nanodrop 2000 (Thermo Fisher Scientific Inc., Waltham, MA, US) in technical triplicates.

#### Cellular lipidome

For extraction, one aliquote in 2 mL Eppendorf vessel, each containing 3 − 10·10^6^ cells were used. 205 µL of cold MeOH (Thermo Fisher Scientific Inc., Waltham, MA, US) were added to the cells, and they were vortexed for 20 s. 10 µL Splash Mix undiluted, 10 µL Ceramide Mix undiluted, and 10 µL Cardiolipin Mix undiluted were added (Avanti Polar Lipids, Alabaster, AL, US). Further, the samples were allowed to thaw at room temperature and were sonicated for 10 min at +4°C. Then, 750 µL of MTBE (Thermo Fisher Scientific Inc., Waltham, MA, US) were added, and the mixture was incubated for 1 h at room temperature under agitation. Phase separation was induced by adding 188 µL water with 0.1% ammonium formate (Thermo Fisher Scientific Inc., Waltham, MA, US). The extract was centrifuged at 10,000 g for 5 min, and two 200 µL aliquots of the upper phase were collected in another 2 mL Eppendorf vessel, dried in a vacuum centrifuge, and stored if necessary at -20°C. For further analyses, samples were dissolved in 200 µL 2-Prop/MeOH/CHCl3 (4:2:1, v/v/v) containing 7.5 mM ammonium formate. LC-MS measurements were performed as described by [61]. A high field Q Exactive HF™ quadrupole-Orbitrap mass spectrometer (Thermo Fisher Scientific) was connected with a robotic nanoflow ion source TriVersa NanoMate® (Advion BioSciences, Ithaca NY, USA) and a nanoelectrospray ionization (nanoESI) chip with spraying nozzles of 5 µm nominal internal diameter. Samples were placed in a 96 twin.tec® well plate (Eppendorf, Hamburg, Germany). Following settings were applied in the Chipsoft 8.3.1 software (Advion BioSciences): ionization voltage 1.25 kV (pos)/ -1.25 kV (neg); backpressure 0.9 psi and in the MS source parameters: capillary temperature +250°C, S-Lens radio frequency level 50. A 9 min polarity switching method with data independent acquisition (DIA) and MS1 scans was used with the following settings: resolution 240,000 (MS1), 60,000 (MS2), AGC target 1e6 (MS1), 2e5 (MS2), maximum IT 150 ms (MS1), 130 (MS2), scan range in MS1 m/z 350-1,050 (pos)/ m/z 200-1,200 (neg), NCE for MS2 of 21 (pos)/ 26 (neg) and a fixed first mass of m/z 80 (pos)/ m/z 150 (neg). LipidXplorer 1.2.8 settings were the following: mass tolerance 5 ppm, min. occupation of 0. In positive mode: intensity threshold 70,000 (MS1)/ 1,000 (MS2), resolution 230,000 (MS1)/ 60,000 (MS2), resolution gradient -180 (MS1)/ -40 (MS2). In Negative mode: intensity threshold 60,000 (MS1)/ 1,000 (MS2), resolution 260,000 (MS1)/ 80,000 (MS2), resolution gradient -170 (MS1)/ -100 (MS2). Identified lipids were removed if the absolute mass error was above 3, if isobaric compounds were present and if the concentration was below LOQ (limit of quantification). LOQ values were calculated by multiplying the standard deviation of 5 repetitive injections of a low concentrated Splash ISTD with 10. For quality control, a 10 µL plasma sample aliquot, which was placed at RT in a 2 mL Eppi, 290 µL Methanol, 10 µL Splash Mix I (undiluted), 1.000 µL MTBE were added. The mixture was shaken for 1 h at RT. Phase separation was induced by adding 250 µL of MS-grade water. Upon 10 min of incubation at room temperature, the sample was centrifuged at 1,000 g for 10 min. The upper (organic) phase was collected in another 2 mL Eppi and dried in the SpeedVac. The dried lipids were dissolved in 200 µL IPA for LC-MS injection.

#### Spent media analysis

##### LC-MS based metabolomics

1 mL suspension was centrifuged (200g, 8 min, +4°C), from which 100 µL liquid media was placed into a 2 mL Eppendorf vessel, and kept on ice. 50 µL of yeast ISTD (extract out of 2 billion cells reconstituted in 2 mL water), and 600 µL methanol (MeOH) were added to reach a final volume of 750 µL (80% MeOH, v/v), which resulted in a 1:7.5 dilution of the sample and a 1:15 dilution of the ISTD. After thorough vortexing, it was kept on ice for 30 min. Then it was vortexed again and kept at -20°C overnight. The samples were centrifuged (14,000g, +4°C, 15 min) and two 200 µL aliquots were dried in a vacuum centrifuge. They were stored if necessary at -20°C. Shipping should be done on dry ice. At least 20·50 µL aliquots of the yeast ISTD (extract out of 2 billion cells reconstituted in 2 mL water) should be placed into 2 mL Eppendorf vessels for internal standardization of the calibrators and QC samples. These samples were also dried in a vacuum centrifuge. All samples were dissolved in 200 µL 20% H2O/80% ACN (v/v). Vigorously vortexed for 3 minutes to ensure complete dissolving of the sample and centrifuged (14,000g, +4°C, 10 min) prior to being transferred to an HPLC vial. LC-MS measurements were performed as described in [61]. A Vanquish™ Horizon HPLC (Thermo Fisher Scientific) with an iHILIC®-(P) Classic (2.1 mm × 100 mm, 5 µm, HILICON, Umea, Sweden) was used for hydrophilic interaction liquid chromatography-high resolution mass spectrometry (HILIC-HRMS). The flow rate was set to 200 µL min-1 and the column temperature to +40°C. Solvent A was 90% 15 mM NH4Ac (pH 9.4), 10% ACN, and solvent B was 90% ACN, 10% 15 mM NH4Ac (pH 9.4). The following gradient was used: 0-12 min ramp from 100% B to 20% B, 12-14 min 20%, 14-17 min ramp to 0% B, 17-19 min 0% B and 20-27 min 100% B as equilibration step. The injection volume was 5 µL and the injector needle was washed with ACN: MeOH: H2O 1:1:1 (v/v/v) for 5 s after each injection. A high field Q Exactive HF™ quadrupole-Orbitrap mass spectrometer (Thermo Fisher Scientific) was used as MS. The following source parameters were applied in both polarities in polarity switching mode: capillary temperature of +280°C, sheath gas flow rate of 40, an auxiliary flow rate of 3, sweep gas of 0, S-lens RF level of 30 and auxiliary gas heater temperature of +320°C applying a spray voltage of 3.5 kV in positive mode and 2.8 kV in negative mode. In MS1 mode the mass range in both polarities was set to m/z 60-900 A maximum IT of 100 ms, a resolution of 120,000, and an AGC target of 3İ0^6^ was applied. Skyline (version 22.2.0.312) was used for peak integration and R/ R studio for final data processing. Identified lipids were removed if the absolute mass error was above 3 and if the concentration was below LOQ. LOQ was defined as the lowest standard concentration in the linear range. For calibration one dried ESTD stock solution is dissolved in 200 µL water for a final conc. of 50 µM. 14 calibrants with concentrations from 25 µM to 0.001 µM were prepared. The internal standard is already dried in the Eppis, therefore the standards are vortexed, centrifuged, and transferred to the well plate.

##### Proline and Glycine

Results for proline and glycine showed inconsistent results. Therefore, these amino acids were quantified using Waters AccQ tagging (Milford, MA, USA) at nine different time points. Briefly, spent media samples were filtered by centrifugation (10 min, RT, 12,000 g) using 3K Amicon (Merck KGaA, Darmstadt, GER) filters to remove proteins. The flowthrough was diluted 1:10 (MilliQ Water) and labeled together with calibration standards according to the manufacturer’s instructions. Norvaline was used as internal standard for normalization. Waters Acquity (UPLC03, Milford, MA, USA) was used as HPLC system. The mobile phase consisted of Eluent A (1:10 diluted in MilliQ Water) and Eluent B (undiluted). For every sample the runtime was set to 11 minutes with a flowrate of 0.7 mL/min. The gradient of Eluent B was increased from 0.1% at the beginning to 90%.

##### Lactate and Ammonia

Lactate and Ammonia were quantified from supernatant for 9 time points. Samples were centrifuged (10 min, 200 g, +4°C) and measured on a BioHT (Roche, Switzerland), following the manufacturer’s instructions. From spent media analyses, limiting compounds were determined, that were fully consumed first. These limiting compounds were then added as supplements to improve the bioprocess

#### Quantification of AAV

##### Droplet digital PCR

As described by [56], for quantifying vector genomes, a droplet digital PCR method from Bio-Rad was employed using the fully automated QX One System (Hercules, CA, USA), which allows for absolute quantification of vector genomes without the need for standard curves. The sample is partitioned into oil droplets, with each droplet serving as an independent PCR reaction compartment. During the initial phase of PCR in the thermal cycler, capsids are degraded, making the DNA accessible for amplification. The vector genome titer is then determined using a droplet reader. To remove extraneous DNA sequences, samples were treated with DNase I (NEB, Ipswich, MA, USA). Prior to this treatment, samples were pre-diluted to enhance the efficiency of DNase I activity. Following droplet generation, PCR was conducted with Bio-Rad ddPCR Supermix (no dUTP) and transgene-specific primers and probe.

##### ELISA

As described by [56], an AAV8 titration ELISA kit (Progen, Heidelberg, GER) was used for quantifying AAV8 capsids, utilizing a monoclonal antibody (ADK8) that recognizes a specific conformational epitope on the assembled AAV8 capsids. The captured AAV 8 particles were identified by the binding of biotinylated anti-AAV 8 ADK8, as this epitope is consistently presented on the AAV 8 capsid structure. Streptavidin peroxidase and a peroxidase substrate were subsequently applied to measure the amount of bound anti-AAV 8, thus indicating the concentration of AAV 8 capsids. The resulting color change was measured at 450 nm using a photometer. The kit also includes an AAV 2/8 particle preparation with a known concentration of labeled AAV 8 particles for calibration purposes.

### Computational Analyses

#### Lipidome analysis

Principal component analysis (PCA) was performed on the lipidomic dataset was performed using the prcomp function in RStudio with the parameters center, and scale set to TRUE. Prior to that, every sample was normalized by dividing each lipid by the total amount of lipid in this sample. Loadings, obtained from PCA, were grouped by lipid class. The five longest loading vectors (lipid classes) were displayed in the PCA plot as well to identify the drivers behind the separation of the samples.

#### Calculation of experimental growth rates during non-exponential growth and production phase

The growth rate was calculated by:

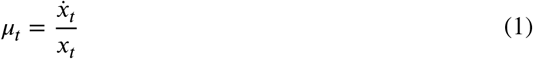

with

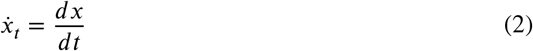

where *μ*_*t*_ is the growth rate at timepoint *t, x*_*t*_ is the biomass value from the fitted function *x*, and 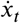 is the derivative of *x*_*t*_. A logistic function *x* was used for fitting, following

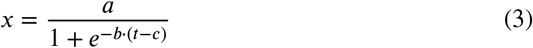

The error of the growth rate was calculated by:

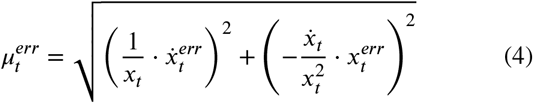

where 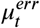 is the error of the growth rate at timepoint *t*, 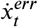 is the error of the derivative value, and 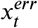 is the error of the fitted biomass value. 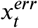 and 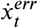 were obtained by:

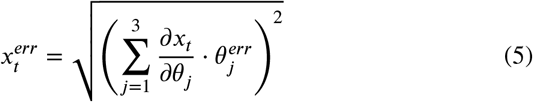

and

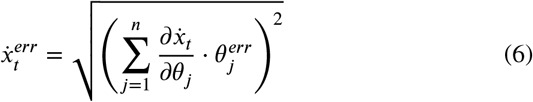

where *θ*_*j*_ is a parameter in the logistic fitting function *x*, and 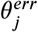 is its corresponding error value from the fitting function *h*.

Derivatives were calculated in R, using the package Deriv (v4.1) [18]

#### Calculation of exchange rates

From the analysis of spent media, we identified 32 metabolites based on their concentration profiles, from which exchange rates could be calculated. These metabolites were categorized into three groups according to the fitting function applied. Specifically, 27 metabolites were fitted using an exponential function, 4 were fitted using a quadratic function, and 1 metabolite was fitted using a cubic function, employing robust fitting techniques (R function nlrob). To determine the exchange rates at each timepoint post-transfection, the derivative of the fitted function was calculated. Error propagation was then performed following the appropriate equation to ensure accurate quantification of exchange rates.

The exchange rate *q*_*i,t*_ of metabolite *i* was calculated as

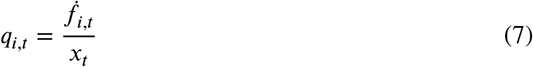

where 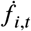 is the derivative value of the fitted function *f*_*i*_ for metabolite *i* at timepoint *t*. The error of *q*_*i,t*_ is calculated by error propagation following:

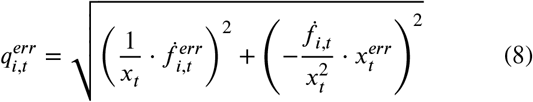

where 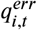 is the error of *q*_*i,t*_ for metabolite *i* at timepoint *t*, 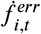 is the error obtained from the derivative 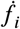 of the fitted function *f*_*i*_. The derivative 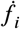 of the fitted function *f*_*i*_ was obtained by

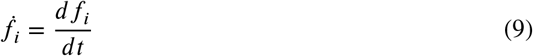

The corresponding error 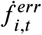 was calculated by:

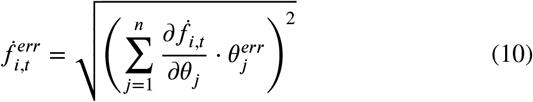

where 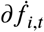 is the partial derivative of the derivative 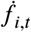 for metabolite *i* at timepoint *t* with respect to the parameter 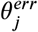 is the obtained error of every parameter *θ*_*j*_ in the fitted function *f*_*i*_ with *n* parameters. To obtain fitted functions *f*_*i*_, the nlrob function in R was used to fit one of the three different functions:

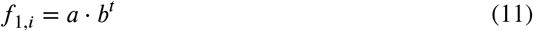

for metabolites, showing an exponential concentration profile,

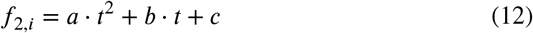

for metabolites, showing a quadratic concentration profile (lactate, proline, and glycine), or,

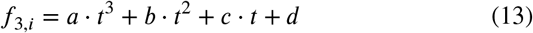

for ammonium, showing a cubic concentration profile.

To consider natural degradation rate of glutamine, media was sampled daily. Glutamine and ammonia were quantified on the BioHT (Roche, Switzerland). The natural degradation rate was calculated using an exponential fit *f*_1,*i*_. The obtained depletion rate was subtracted from the derivative value 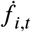 for glutamine or added to the derivative value 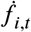 for ammonia.

#### Reconstruction of context-specific models

To reconstruct context-specific GSMMs, the most recent GSMM of the human metabolism, human1 [59], was used. Initially, the uptakes of 42 compounds were identified to enable feasible solutions of the GSMM. These compounds were selected based on a chemically defined medium (22400 - RPMI 1640, HEPES, Thermo Fisher Scientific Inc., Waltham, MA, USA), since the exact composition of the used F17 expression media is currently not available. Additionally, all compounds identified by [22] that are not essential for growth were included as potential secretion reactions, totaling 23 compounds. Moreover, metabolites for which exchange rates could be calculated (as detailed in the subsequent section) were incorporated into the model as constraints. The compounds in the growth medium were assigned exchange reaction boundaries of [-1,000, 0], while the boundaries for all other exchange reactions were set to [0, 1,000].

To obtain context-specific GSMMs, transcriptomic data from the exponential growth phase were utilized as reported in [23]. This transcriptomic dataset, derived from a non-producing strain, encompasses four time points during exponential growth, with transcriptomes sequenced in triplicates. For each sample, a GSMM was reconstructed using the CORDA algorithm (v 0.4.2) [62]. To ensure the preservation of conserved reactions during reconstruction, five subsets of each transcriptome were generated. These subsets consisted of percentiles of the top expressed genes (10%, 25%, 50%, 75%, and 90%). Corresponding reactions from each subset of genes were extracted based on gene-reaction rules and assigned a confidence score of 3, ensuring the highest score in CORDA. Additionally, the biomass reaction, reactions for compounds from the growth medium, and potentially secreted reactions were also assigned a confidence score of 3. All remaining reactions were assigned a confidence score of -1, following the assumption that the genes were not expressed. Reactions from all reconstructions were combined to create a context-specific GSMM. This “adapted” CORDA approach was compared to the default method [62], which involved assigning confidence scores ranging from -1 to 3 to reactions based on gene expression levels. Validation was conducted by integrating provided exchange rates, calculated from the same samples [23]. Predicted growth rates were obtained from flux balance analysis (FBA), using Cobrapy (v 0.22.1) compared to experimental growth rates [23] and to predicted growth rates from the same process using a manually curated GSMM [58], based on Recon2 [66].

Next, we utilized recently published transcriptomic data from the identical process described in [56] to reconstruct 80 context-specific GSMMs at time points 0, 4, 24, 48, and 72 HPT. During the subsequent optimization of production reactions, it was noted that the transport reactions of three nucleotides from the cytoplasm into the nucleus were not included in the models and were therefore added manually. To validate the reconstruction, we employed two approaches: first, we applied logisitic principal component analysis (LPCA) [41] to the differential reaction set [82] of all models and compared the results to those obtained from PCA applied to transcriptomes as shown in [56]. Secondly, we integrated the calculated exchange rates and obtained simulated growth rates using FBA, where the lower bound (*lb*_*i*_), and upper bound (*ub*_*i*_) for each metabolite *i* was calculated by:

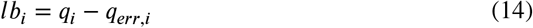

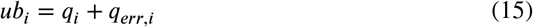

These simulated growth rates were then compared to experimentally determined growth rates. Production reactions for AAV were incorporated into the models using Biopython (v1.79) and AAV genomes (NCBI UID: 5075992). After integrating the production reactions into the models, we analyzed the production envelope at each timepoint and performed comparative analyses between the cell lines.

#### Calculation of specific production rate from experimental titers

First, production reactions for AAV production were added. The viral genome from AAV 8 was downloaded and the sequence for viral proteins VP1, VP2, and VP3 was extracted. The ratio of 1:1:10 (VP1:VP2:VP3) was considered in the production reaction [60]. Additionally, the replicated genome was considered as 4.8 kb long and consisting of 25% Adenine, 25% Guanine, 25% Thymine, and 25% Cytosine. Moreover, we added transport reactions for all amino acids into the nucleus, since AAV production occurs in the nucleus [52]

- Production of VP1 in the nucleus
- Production of VP2 in the nucleus
- Production of VP3 in the nucleus
- Assembly of a full capsid with ratio 1:1:10
- Production of transgene
- Assembly of full capsids
- Transport of full capsids from nucleus into the cytoplasm
- Transport of full capsid from cytoplasm outside of the cell
- Exchange reaction for full capsid

See supplemental material for a detailed explanation of the added reactions for AAV production. Additionally, we added transport reactions of amino acids to the nucleus, since AAV replication happens in the nucleus [52].

We calculated specific production rates from experimental data. Experimentally, titers could only be determined at 24, 48, and 72 HPT, as the productivity at 4 HPT was too low to measure accurately. Consequently, we calculated the percentage of biomass allocated to production at 24, 48, and 72 HPT to minimize discrepancies with the experimentally determined titers. This was done by setting the lower bound (*lb*) of the biomass reaction to the maximum growth rate, multiplied by a specific value *α*_*t*_ for each strain and time point.

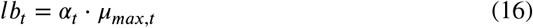

After that, *α* was determined by optimizing the production reaction, so that the squared difference between the simulated production rate and the experimental production rate is minimal. For optimization, we used a `differential_evolution` algorithm from scipy (v 1.7.3) with bounds between 0, and 1, since the *α* values should describe a percentage of the maximal biomass, that is used for production of AAV. We used the `best1bin` strategy with a maximum of 10 iterations, a population size of 15, and a tolerance of 10·10^−7^. Using these factors (*α*), we fitted a quadratic function, also assuming that at time point 0 HPT, all resources are directed towards growth giving an *α* value of 1, to predict an *α* value at 4 HPT. This predicted *α* value was then used to calculate the specific production rate in the GSMMs at 4 HPT. All obtained *α* values (percent allocated to production and percent to biomass) were plotted along with their fitting functions. After determining all production rates, we generated a realistic production envelope to compare the two cell lines by fixing simulated growth rates and maximizing production rates.

#### Parsimonious FBA for pathway analysis

For all context-specific GSMMs, we performed parsimonious flux balance analysis (pFBA). The resulting flux distribution was normalized by dividing by the simulated growth rate at each time point and state. Individual cell line-specific GSMMs were compared in a pairwise manner. Reactions that consistently exhibited more than 100% higher or lower flux across all time points, or over the first three time points, between the two strains were identified and analyzed.

### Fermentation for validation

Fermentation was repeated using 1 mg HIF-1*α* inhibitor PX-478 dihydrochloride (Merck KGaA, Darmstadt, Germany), dissolved in 1 mL F17 media per 250 mL fermentation suspension. Additionally, limiting amino acids, aspartate, and glutamate (Merck KGaA, Darmstadt, Germany) were added as an alternative to improve the bioprocess (1.5x amount of initial amount, 0 HPT). Control settings were used as described before (see section Bioprocess and Sampling).

### Code and data availability

All generated, and used data, as well as analyses are provided on GitHub: *github.com/LeoZ93/hek293aav*.

## Results and Discussion

### HP cells are larger but lighter than LP cells

The dry cell mass and the composition of biomass– including total proteins, lipids, DNA, RNA, and carbohydrates– were quantified at five time points during fermentation for both the HP and LP strains. In both strains, transfection had no influence on total mass or composition. However, the dry mass decreased over time, with the LP strain showing a steeper decline.

As shown in Fig 2a, the dry mass of the HP strain ranged from 295 pg/cell to 380 pg/cell, with an average of 339.2 pg/cell, demonstrating more consistency over time compared to the LP strain, which ranged from 298 pg/cell to 447 pg/cell, averaging 388.9 pg/cell (adjusted p-value 6.515·10^−4^, t-test, Bonferroni correction).

**Figure 2:**
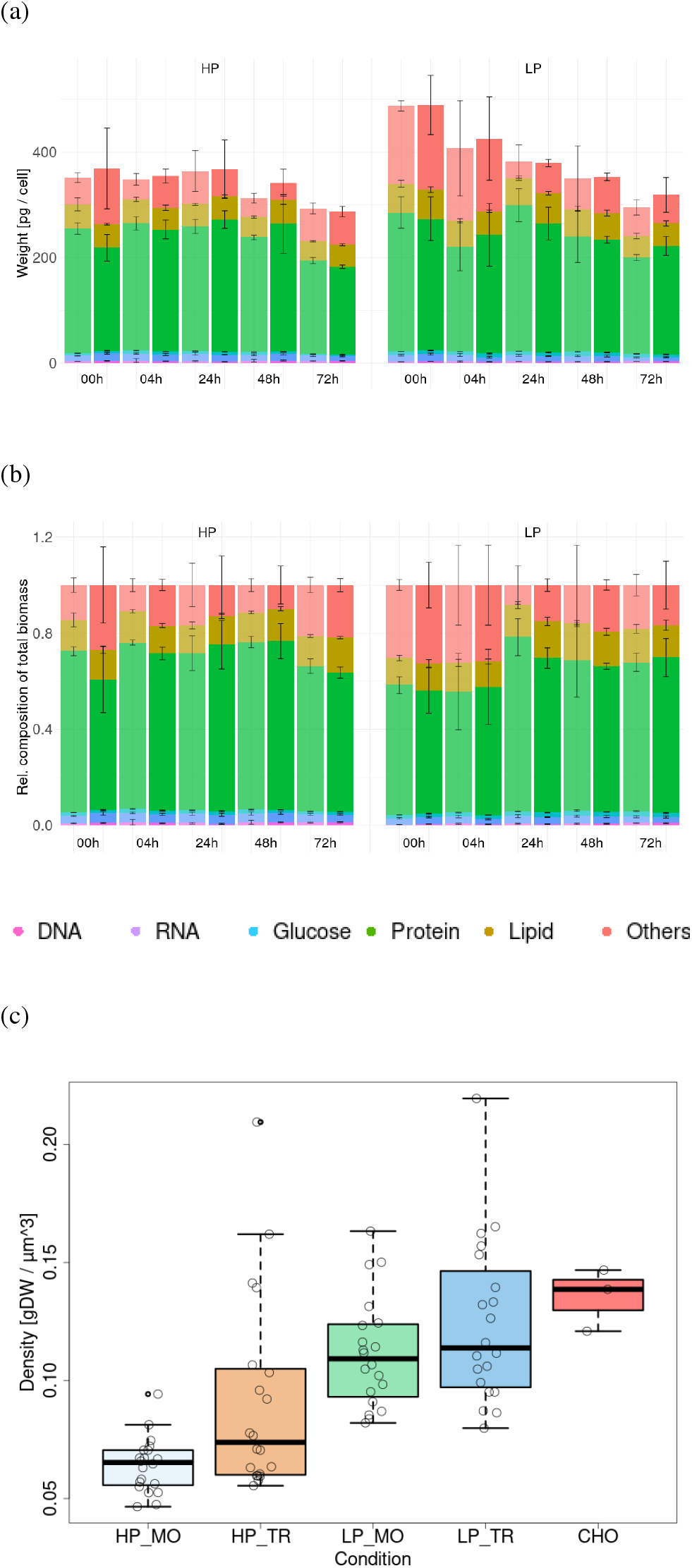
Absolute (a) and relative (b) biomass composition, and volumetric mass density (c) for HP and LP strains. The LP had a higher dry mass than the HP (adjusted p-value 6.515·10^−4^, t-test, Bonferroni correction) and density (adjusted p-value 6.805.10^−7^, t-test, Bonferroni correction), but a larger volume (adjusted p-value 8.637·10^−7^, t-test, Bonferroni correction). In the first 4 h, the HP had a higher relative protein content, while other components were similar between strains. Transfected states are displayed fully colored, while mock states are transparent.

For both strains, this was lower than the previously reported 514 pg/cell [23, 58]. However, this difference may be attributed to variations in cell composition, as the data in [58] were derived from multiple human cell lines based on Recon2 [68]. In comparison, the average biomass for CHO cells is 264 pg/cell [67], making the dry mass in our study generally higher.

The relative composition analysis showed that the contributions of lipids, DNA, and RNA remained almost constant over time and were independent of the strain (Fig 2b). In contrast, the HP strain had a higher protein content compared to the LP strain, particularly during the first 4 HPT. Since the amounts of lipids, DNA, and RNA were similar in both strains, the higher protein content in the HP strain suggests that other components contributing to dry mass must be higher in the LP strain. In fact, 35% of the dry mass in the LP strain, especially early post-transfection, remains unaccounted for. A similar high proportion of unknown dry mass has also been observed in some CHO cells [67].

The HP strain displayed a higher volume regardless of the production state of the strain, despite having a lower dry mass compared to the LP strain (Fig 2c).

### HP stores energy as triacylglyceride; LP prefers glycogen

Next, we investigate the lipid profile for both strains that on average represented approximately 10% to 18% of the total biomass irrespective of the strain. We observed that phosphatidylcholine (PC) and phosphatidylethanolamine (PE) dominated across all time points (Fig 3b) accounting for about 65% of the total lipids and reflecting their essential role in mammalian cell membranes [20, 70]. Our distribution aligns with previous analyses of HEK293 strains during exponential and stationary phases [83] and mirrors typical mammalian cell lipid profiles [72, 73].

**Figure 3:**
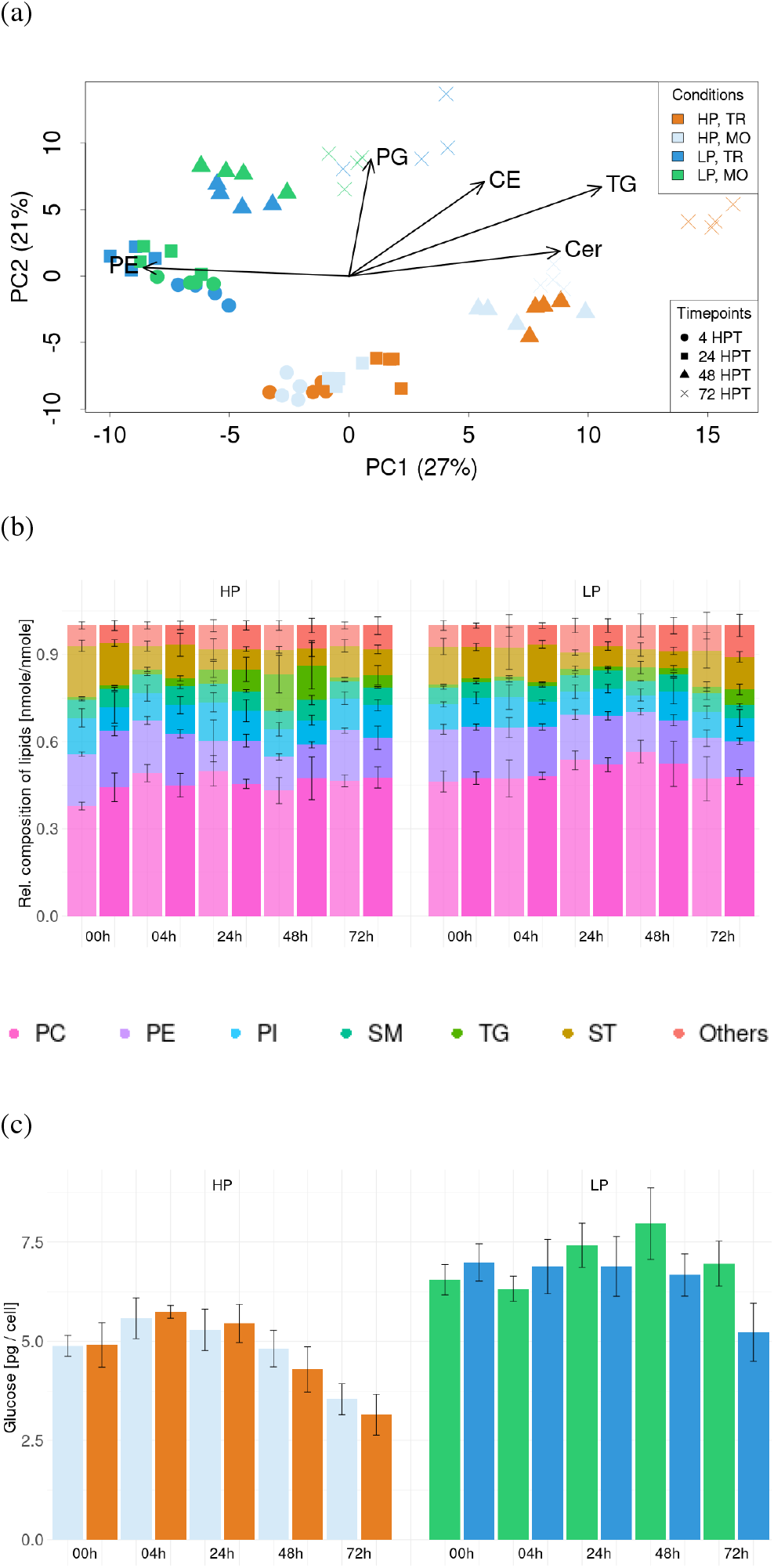
PCA of lipid profiles (a), relative lipid (b), and glucose (c) composition of HEK293 strains during AAV 8 production and mock states. PCA resulted in a clear separation of HP and LP strains (a), with loading values grouped by the five most important groups. The predominant drivers for the separation of the strains were TG, and Cer, pinpointing in the direction of the HP cluster, while PR and PG loading vectors directed in the LP cluster. PE, PI, and PC constitute the majority of lipids in both, HP and LP strains. TG amounts changed most significantly from 1% to 12% in the HP strain until 48 HPT, indicating that the HP strain stored glucose as TG (b), while the LP strain stored glucose predominantly as glucogen (c).

PCA analysis clearly distinguishes between HP and LP, regardless of whether the cells were mock-transfected or transfected (Fig 3a). This separation is mainly driven by differences in triacylglycerides (TG), ceramide (Cer), phosphatidylethanolamine (PE), and phosphatidylglycerols (PG). HP cells show higher levels of TG and Cer but lower levels of PE and PG compared to LP. These lipid concentrations also change over time, with TG and Cer increasing as fermentation progresses, while PE decreases (Fig S1). In the HP strain, phospholipid levels ranged from 5% to 12% (Cer), 7% to 13% (PG), and 10% to 19% (PE), whereas in the LP strain, they ranged from 4% to 10% (Cer), 7% to 18% (PG), and 12% to 18% (PE).

The greatest difference was observed in TG levels, which rose from 1% to 12% in HP during the first 48 h, compared to only 1% to 5% in LP. Interestingly, the LP strain appeared to store glucose as glycogen, as indicated by its higher glucose content relative to HP (Fig 3c), suggesting a potential metabolic bottleneck in LP, possibly due to oxygen limitation (see below), since glycogen is a less efficient form of energy storage compared to fatty acids [6]. Additionally, HP cells displayed more dynamic changes in storage capabilities over time compared to LP, further highlighting differences in energy storage strategies between the strains.

Cholesterol esters (CE) increased similarly in both strains, correlating with the progression of fermentation rather than distinguishing between them.

Finally, we speculate that lipid remodeling occurs following transfection, characterized by an initial increase in short-chain fatty acids shortly after transfection, with the lipid profile gradually returning to its original distribution by the end of fermentation (Fig 3b). However, this effect is less pronounced and appears less evident in the LP strain.

### GSMMs capture transcriptomic and growth trends

To further investigate the metabolic differences between HP and LP, we reconstructed strain-specific GSMMs using CORDA [62]; see Methods for details. These models are based on the integration of the biomass composition (reported above), together with transcriptome and exometabolome measurements collected throughout the fermentation process.

To validate our reconstruction pipeline, we used transcriptomic data and exchange rates from independent studies [23, 58] to build a strain-specific GSMM for a non-producing HEK293F strain. The model predicts a specific growth rate of 0.0301 h^−1^, which closely matches the experimentally measured rate of 0.0281 h^−1^, resulting in a relative error of 7.1% (Tab 2). This solid agreement indicates a good accuracy of our model and validates our reconstruction approach.

**Table 2.**
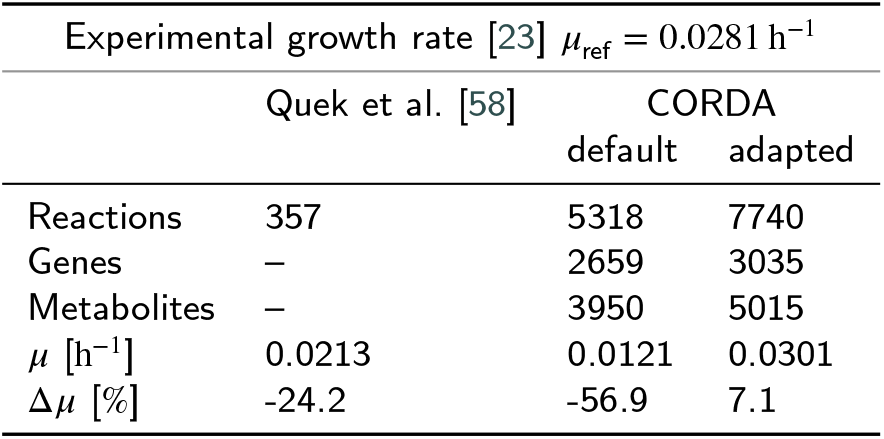
Comparison of the number of reactions in HEK293F-specific GSMMs and their predicted growth rates (*μ*) with the measured exponential growth rate (*μ*_ref_) as reported in [23]. ‘Default’ and ‘adapted’ CORDA [62] refer to the models developed in this study (see methods); Quek et al. [58] represents a previously established model. The relative error Δ*μ* is defined as *μ/μ*_ref_ − 1.

| | Experimental growth rate [23] $\mu_{\text{ref}} = 0.0281 \text{ h}^{-1}$ | | |
| --- | --- | --- | --- |
|  | Quek et al. [58] | CORDA default | CORDA adapted |
| Reactions | 357 | 5318 | 7740 |
| Genes | — | 2659 | 3035 |
| Metabolites | — | 3950 | 5015 |
| $\mu \text{ [h}^{-1}\text{]}$ | 0.0213 | 0.0121 | 0.0301 |
| $\Delta\mu \text{ [\%]}$ | -24.2 | -56.9 | 7.1 |

Next, we reconstructed 80 context-specific GSMMs for each of the four replicates of our two producer strains at five time points, using transcriptomic data from Pistek et al. [56], and the corresponding media composition. To assess whether these models captured the differential information present in the transcriptome (Fig 4c), we collected model metrices (Tab 3), and applied LPCA [82], a method we previously demonstrated to replicate clustering patterns observed in transcriptome-based PCA using only GSMMs. As expected, LPCA of the reconstructed metabolic models revealed clustering patterns similar to those from transcriptome-based PCA (compare Fig 4c with Fig 4b). These findings suggest that even at the metabolic level strain differences were more pronounced than between conditions or time points.

**Table 3.**
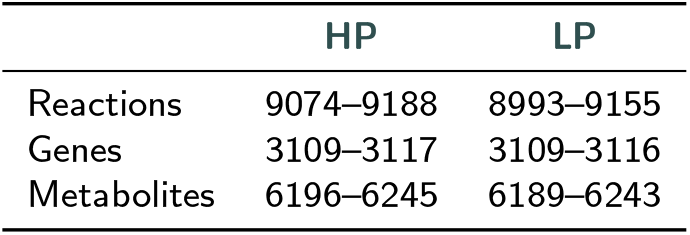
Comparison of Reactions, Genes, and Metabolites in HEK293-specific GSMMs. Numbers were extracted at four time points (4 HPT to 72 HPT) for MO and TR. Values represent minima and maxima across all time points and conditions.

**Figure 4:**
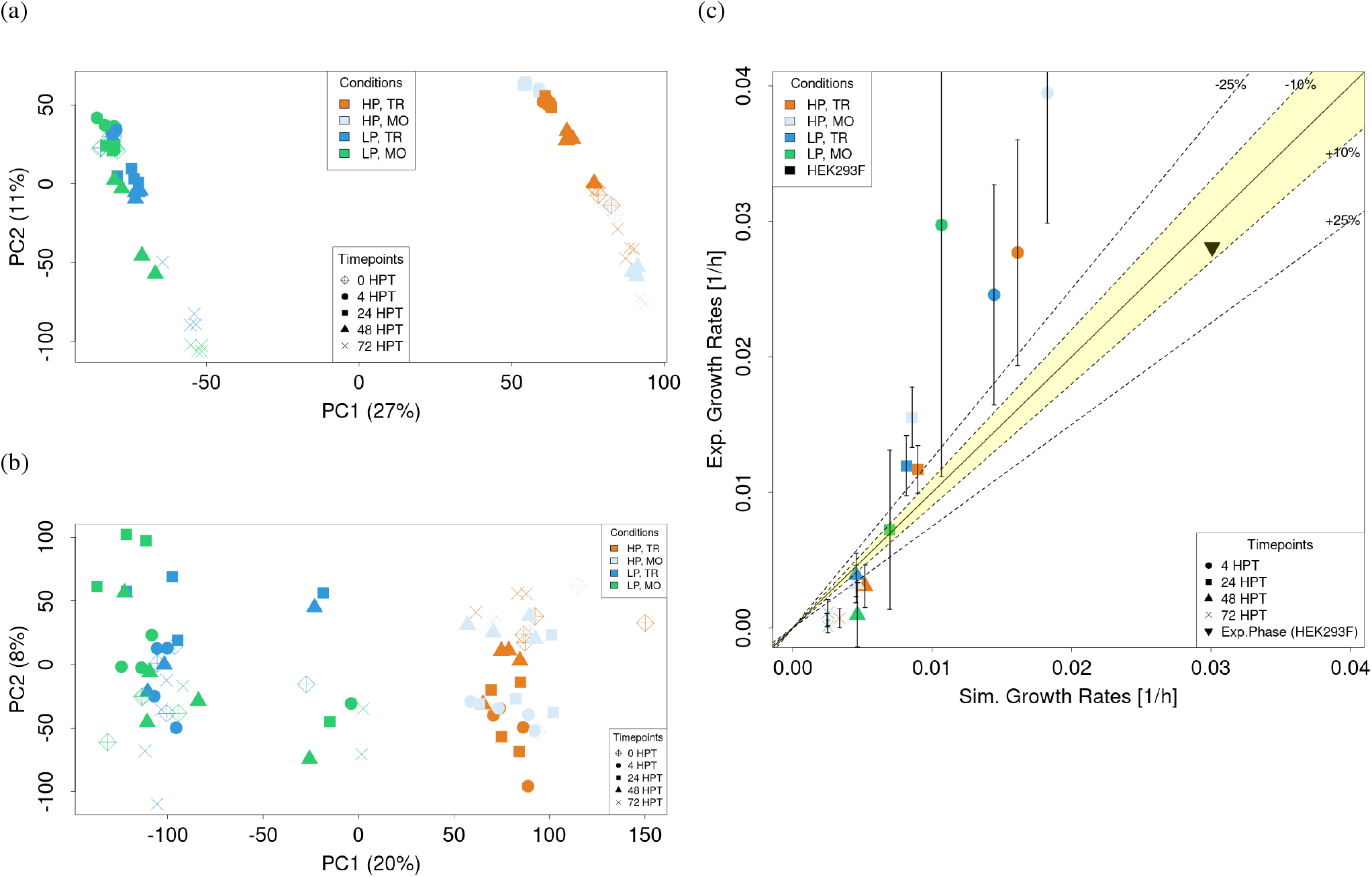
PCA from transcriptomic data (a), LPCA of binary reactions matrix derived from GSMMs (b), and simulated vs. experimental growth rates (c). When comparing LPCA (b) with a PCA plot performed on whole transcriptomes (a) we observed a similar clustering, indicating that GSMMs capture differences between strains in a similar way. Growth rates were calculated from dry mass measurements. Error bars were obtained after fitting the regression function (see methods). Simulated growth rates were obtained after merging biological replicates. Aside from the first time points 4 HPT, the simulated and experimental growth rates are comparable, indicating that the reconstructed GSMMs can capture realistic growth rates during AAV production and in mock state (c).

Finally, to assess the quality of our context-specific GSMMs for both producer strains, we used FBA to simulate growth rates over 4 HPT to 72 HPT, using the measured exchange spectrum as input, and compared the results to a logistic fit of the experimentally measured growth curve.

We observed reasonable agreement (±25 %) at intermediate growth rates, while lower rates were overestimated (Fig 4a). For higher growth rates, the large deviations were possibly due to reduced reliability in the experimental data, as indicated by substantial error bars. Nevertheless, the error bars of most experimental data points at least intersect with the ±25% region around our predictions (Fig 4a), supporting the overall robustness of our model predictions.

### HEK293 strains differ in biomass composition and limiting compounds in spent media

In addition to biomass analyses, we performed a spent media analysis as described in the methods. We identified the amino acids aspartic acid in the LP strain and serine in the HP strain as limiting, regardless of the state (see Fig S3–Fig S7).

### HEK293 strains have similar AAV production potential, but drastically lower actual rates

After integrating AAV particle production reactions into the GSMMs of the HP and LP strains (see Supplementals), we applied FBA to compute production envelopes at four time points ranging from 4 to 72 HPT (Fig 5). These envelopes represent the maximum production capacity of each strain as a function of growth rate. We found that, for the experimentally measured uptake rates, both strains exhibit nearly identical production capabilities. This means that despite the differences in their GSMMs, the overall production potential of HP and LP is stoichiometrically almost the same. However, while the actual production rate in HP is up to six times higher than in LP, both strains produce at rates that are two orders of magnitude lower than their theoretical maximum production capacity.

**Figure 5:**
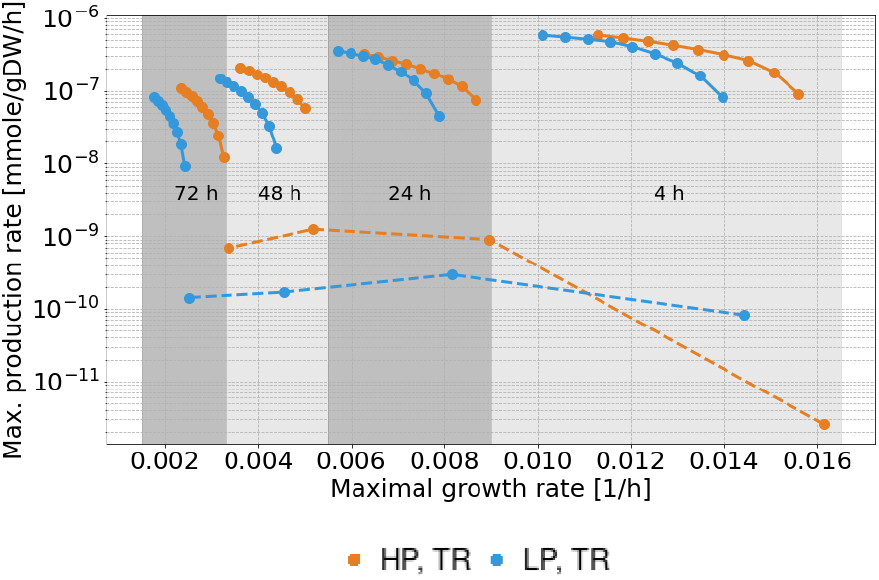
Production envelope of HEK293 strains (HP: orange, LP: blue) 4 HPT, 24 HPT, 48 HPT, 72 HPT (solid lines), as well as the real production envelope (dashed line). For every time point and cell line, the maximum growth rate was determined by FBA. These maximum growth rates were then fractionally reduced to 70% and fixed to the *lb*_*bm*_, before optimizing for AAV production. The resulting solid lines displayed at the top of the plot represent the maximum production capacity. The production envelope based on experimental data (dashed line) was obtained after calculating the fraction of biomass, used for production. While the experimental production rate of the HP strain is higher compared to the LP strain, both strains are two orders of magnitude lower than their theoretical maximum production rate.

### Reaction analysis of HEK293 context-specific GSMMs during AAV 8 production identifies pseudohypoxia in the LP strain

To better understand why the LP strain performs poorly compared to the HP strain and to identify potential targets for improving LP productivity, we performed pFBA for both transfected (TR) and mock (MO) conditions.

In the TR models, growth rates were fixed to the simulated values *α*_*t*_*μ*_*sim,t*_ (Fig S2), and production reactions during AAV production were optimized (Fig 5). For the MO models, the biomass reaction was optimized. This allowed us to compare flux distributions across both strains and conditions. To identify significant differences, we classified reactions as differential if their flux values consistently varied by at least 50% between strains across the first three time points (4 HPT to 48 HPT) or across all time points. These early time points were selected since the onset of cellular responses to viral infection typically occurs within 1–4 HPT [33].

Through this analysis, we identified 40 differential reactions: 18 exhibited higher flux in the LP strain (mock: 5, transfected: 13), and 22 showed lower flux compared to the HP strain (mock: 7, transfected: 13) (see Tab 4). Out of these, 28 reactions had gene associations, and 12 were exchange reactions.

**Table 4.** Comparison of reactions with consistently higher (red) or lower (blue) fluxes in the GSMMs of LP vs. HP over the first three time points (4 HPT to 48 HPT). Each reaction is listed with its gene-protein-reaction (GPR) mapping, where overexpressed genes are in red and underexpressed genes in blue. Biochemical functions and metabolic subsystems are also included. Bold genes indicate regulatory targets of HIF-1*α* (see main text).

| Reaction ID | Gene-Reaction Rule | Reaction |  | Subsystem |
| --- | --- | --- | --- | --- |
| MAR09061 | – | ala[e] | → | Exchange reactions |
| MAR09068 | – | pro[e] | ← | Exchange reactions |
| MAR09071 | – | glu[e] | ← | Exchange reactions |
| MAR09685 | – | sbt [e] | → | Exchange reactions |
| MAR09012 | – | 2-oxo–3-methylvalerate [e] | → | Exchange reactions |
| MAR09001 | – | 2-oxo–3-methylvalerate [e] | ← 2-oxo–3-methylvalerate [c] | Transport reactions |
| MAR05330 | **SLC1A3** or *SLC1(A1 or A2 or A6 or A7) or SLC6A19 or SLC25A20* | Na<sup>+</sup> [e] + glu[e] | → Na<sup>+</sup> [c] + glu[c] | Transport reactions |
| MAR04312 | **COPB2** or *ARCN1 or COPB1 or COPE or COP(G1 or G2)* | sbt[e] | → sbt[c] | Transport reactions |
| MAR05481 | **SLC7A5** | ala[c] + pro[e] | → ala[e] + pro[c] | Transport reactions |
| MAR05517 | **SLC7A5** | met[e] + pro[c] | ← met[c] + pro[e] | Transport reactions |
| MAR09031 | **SLC22A7** | glu[c] + orot[e] | → glu[e] + orot[c] | Transport reactions |
| MAR04875 | – | orot[c] | → orot[e] | Transport reactions |
| MAR08992 | **RPE** | 2 ru5p[c] | → 2 xu5p[c] | Pentose phosphate pathway |
| MAR04404 | **TKTL1** or *TKTL2 or TKT* | xu5p[c] + e4p[c] | → GAP[c] + f6p[c] | Pentose phosphate pathway |
| MAR04501 | **TKTL1** or *TKTL2 or TKT* | xu5p[c] + r5p[c] | → GAP[c] + s7p | Pentose phosphate pathway |
| MAR04565 | **TALDO1** | GAP[c] + s7p[c] | → e4p[c] + f6p[c] | Pentose phosphate pathway |
| MAR04304 | **G6PD** | NADP<sup>+</sup> [r] + g6p[r] | → H<sup>+</sup> [r] + NADPH[r] + 6 pgl[r] | Pentose phosphate pathway |
| MAR00710 | **IDH1** or *IDH2* | NADP<sup>+</sup> [c] + icit[c] | → AKG[c] + CO<sub>2</sub> [c] + NADPH[c] | TCA cycle |
| MAR03806 | **ALDH4A1** | H<sub>2</sub>O[m] + NAD<sup>+</sup> [m] + glu5sa[m] | → 2 H<sup>+</sup> [m] + NADH[m] + glu[m] | Arg and pro metabolism |
| MAR05127 | – | H<sup>+</sup> [c] + NH<sub>3</sub> [c] | ← NH<sub>4</sub><sup>+</sup> [c] | Miscellaneous |

#### Consistent metabolic changes

- **Altered amino acid Exchange**. In the LP, alanine, glutamate, and proline uptake were consistently higher across both mock and transfected states, while glutamine secretion was reduced. In contrast to these flux differences, the gene *SLC7A5*, part of the SLC transporter family, had significantly lower expression in the LP compared to the HP across all time points and conditions (p-value < 3.04·10^−10^, t-test).
- **Upregulation in PPP**. The LP showed higher activity in the PPP, a pathway often up-regulated in cancers to support rapid proliferation. Specifically, transketolase (*TKTL1*) were more active, suggesting a metabolic shift toward enhanced nucleotide synthesis and improved redox balance.
- **Downregulation in tricarboxylic acid (TCA) cycle**. In the LP, downregulation of isocitrate dehydrogenase (*IDH1*) reduces the conversion of isocitrate to *α*-ketoglutarate. This affects NADPH production and reduces flux through the TCA, indicating a shift away from oxidative metabolism.

These metabolic changes resemble hallmarks of cancer metabolism [54]. Since some of these responses are (at least partially) mediated by hypoxia-inducible factors (HIFs), we explored the potential association with HIF-1*α*, though other mechanisms could also contribute.

In fact we find, *SLC7A5* is often upregulated in tumor environments under hypoxia [50, 26] consistent with increased amino acid uptake flux in the LP. *TKTL1* expression is induced by hypoxia mediated by HIF-1*α* in colorectal cancer [8]. Aberrant *TALDO1* activity is implicated in various malignancies, suggesting its involvement in metabolic reprogramming that supports rapid cancer cell growth and survival [77].

Based on our observations, we hypothesize that HIF-1*α* is activated in the LP strain despite adequate oxygen levels. This suggests a state of pseudohypoxia, where hypoxia-like conditions are mimicked even with sufficient oxygen [31]. In fact, HIF-1*α* expression was significantly higher in the LP strain compared to the HP strain across all time points and conditions (adjusted p-value < 2.93.10^−17^, t-test, Bonferroni correction). This phenomenon could be driven by factors such as high glucose uptake, mitochondrial dysfunction, and an energy imbalance favoring NADH, similar to observations in HEK293 strains adapted to suspension culture [36].

### Experimental validation

Based on our hypothesis, we inhibited the transcription factor HIF-1*α* using PX-478 dihydrochloride, a known HIF-1*α* inhibitor [76, 40, 84, 63, 78]. As expected, PX-478 completely inhibited cell growth in the LP strain, confirming its strong dependence on HIF-1*α* (Fig 6a, light blue). In contrast, adding aspartic and glutamic acid (Fig 6a, dark red), identified as limiting compounds in the spent media, did not significantly impact cell growth compared to the control fermentation (Fig 6a, dark blue).

**Figure 6:**
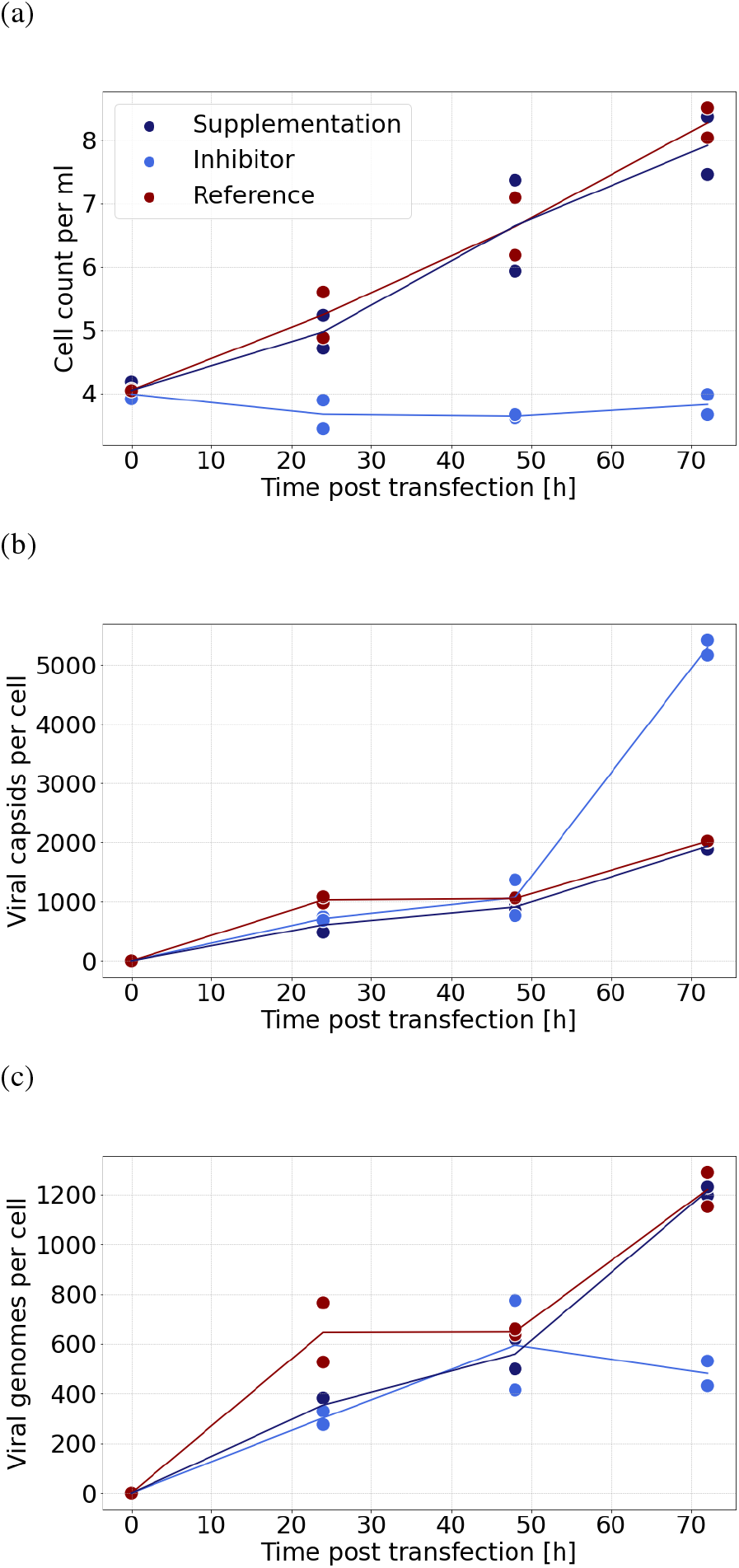
Validation experiment, when inhibiting HIF-1*α* in LP HEK293 strain during AAV production. HIF-1*α* inhibition (light blue), showed halted growth (a), higher viral capsid production (b), but reduced viral genome production (c), while the production under original conditions (dark blue), and with aspartic acid, and glutamic acid supplemented (dark red), resulted in similar growth rates (a), and production rates (b, c).

Next, we assessed productivity under these conditions by measuring viral capsid concentration (Fig 6b) and viral genome concentration (Fig 6c). Interestingly, inhibiting HIF-1*α* nearly doubled per-cell capsid production compared to the control (p-value < 0.006727, t-test), but viral genome production decreased. This decline might be due to reduced activity in the PPP, which is tied to HIF-1*α* and vital for nucleotide synthesis. However, since AAV genome replication depends on human DNA replication and repair proteins (e.g., MCM, Polymerase delta) [51], inhibiting HIF-1*α* may decouple PPP from HIF-1*α* regulation, possibly improving viral genome replication efficiency.

## Conclusion

HEK293 strains are currently the most prominent platform for the production of AAV particles in gene therapy [12]. Despite numerous advances in the past decades [19, 65, 9], production titers remain insufficient to meet necessary demands [52]. In this study, we present a comprehensive multiomics analysis, encompassing transcriptomic, lipidomic, and fluxomic data, to investigate and compare two different HEK293 strains in both mock and AAV production states.

When comparing the HP HEK293 strain with the LP strain, we found that the most significant differences were between the strains themselves rather than between their production states, corroborating findings by [56]. Recent studies on HEK293 cells have revealed high glucose consumption and elevated production of lactate and succinate [36], indicative of pseudohypoxia in HEK293 cells recently adapted to suspension culture. These observations were reproduced in subsequent studies during AAV production, which also showed upregulated immune responses [56] and unfolded protein responses [43]. Both of these responses have been associated with HIF-1*α* activity in solid tumor environments [5, 24]. By employing a multi-omics approach and integrating GSMMs, we identified that the LP strain experiences activated HIF-1*α* due to pseudohypoxia. Lipidomic analysis showed higher concentrations of ceramides in HP strains, indicating a tumor-like state in the LP strain. Exometabolomic analysis revealed that the LP strain preferentially consumes aspartate and glutamate, a common behavior of solid tumor cells to maintain energy balance [28]. Additionally, we identified 17 reactions consistently upregulated in the GSMMs derived from the LP strain compared to the HP strain, with 15 of these linked to HIF-1*α* activity or hypoxic environments. After identifying HIF-1*α* as a potential target, we observed that it was significantly upregulated in the LP strain compared to the HP strain (adjusted p-value < 2.931·10^−17^, t-test, Bonferroni correction). This identification would have been more difficult in whole transcriptome analysis and GSEA as performed by [39, 56]. Inhibiting HIF-1*α* immediately halted cellular growth in the LP strain and doubled AAV capsid productivity. However, the number of viral genomes per cell decreased compared to standard production in the LP strain, likely due to insufficient nucleotide synthesis through the PPP. The PPP is central to synthesizing precursors for *de novo* nucleotide synthesis and is closely tied to HIF-1*α* activity in cancer cells. Inhibiting HIF-1*α* likely downregulated *de novo* nucleotide synthesis, leading to fewer viral genomes. To mitigate this bottleneck, additional nucleotides could be supplemented in the media to maintain viral genome production. The overexpression of HIF-1*α* has been exploited in bioprocesses involving CHO cells to enhance RP production. Specifically, the target gene was placed under the control of a HIF-1*α*-responsive sequence, thereby ensuring that the RP is expressed upon HIF-1*α* activation in CHO cells. This activation was achieved by inducing hypoxia through reduced oxygen flow [80]. These findings suggest that hypoxia and the activation of HIF-1*α* may serve as critical regulators for bioprocess control and AAV production.

Aside from these results, the optimization of a GMP-conform bioprocess should not include any substances or additives, such as inhibitors, due to potential difficulties in downstream purification, which was not considered in this study. Therefore, it is necessary to create stable knockout strains, to inhibit the function of HIF-1*α*, instead of using inhibitors, such as PX-478.

## Acknowledgements

The authors kindly thank the whole GTPD Upstream Team 1, specifically Manfred Ostermann, Katharina Portner, Ronald Leitner, Sonja Muellner, Robert Pachlinger, Susi Heider, Carolin Kahlig, and Lucia Micutkova, as well as Peter Andorfer, Peter Eisenhut, Bernd Innthaler, Felix Fuchsberger, Fabian Knofl, Andrea Seybel, Sabine Unterthurner, and Robert Pletzenauer for their support.

## Supplementary Information

**Figure S1:**
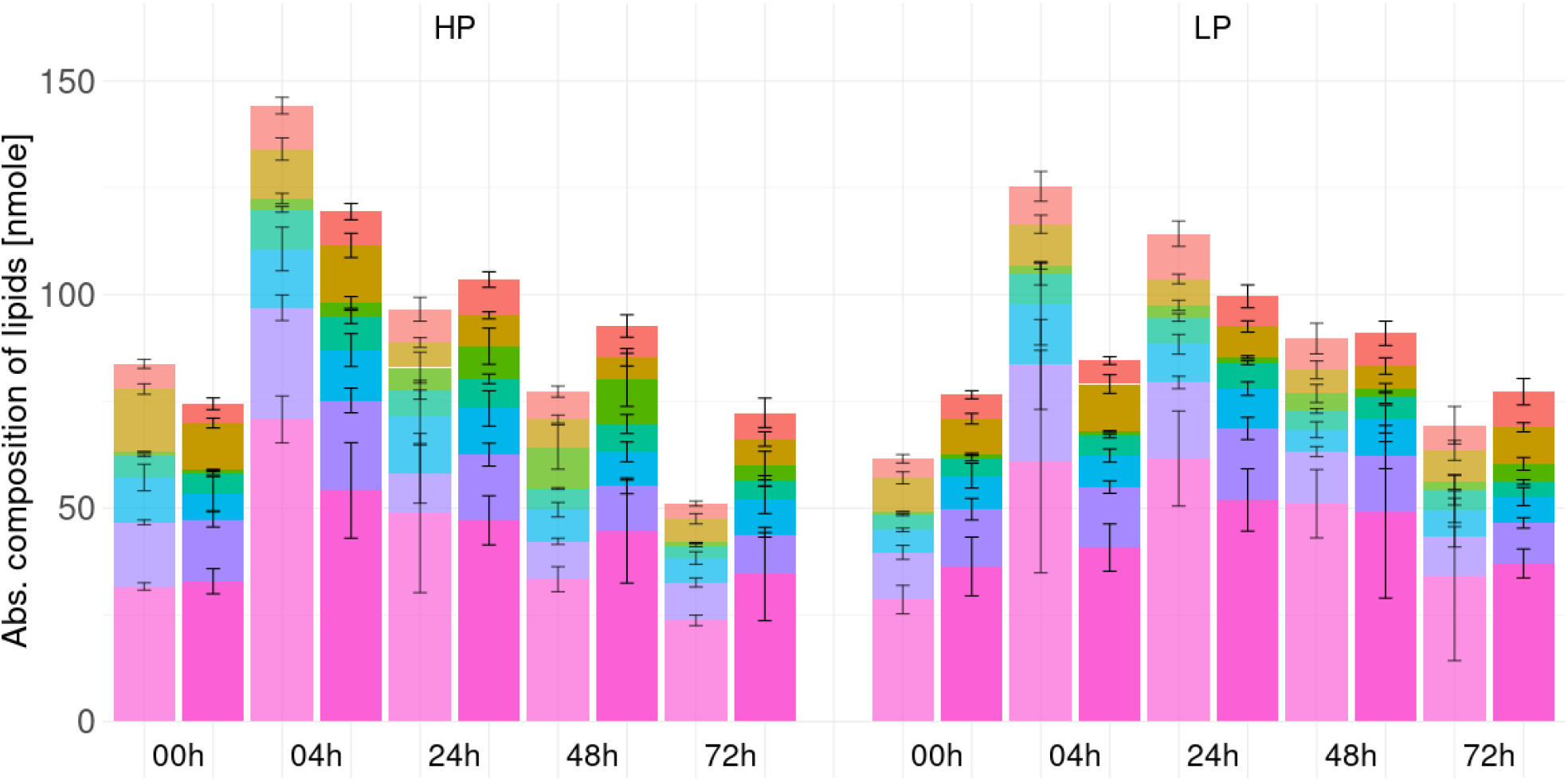
Absolute lipid composition of HP and LP strains during AAV 8 production and during growth. Lipid profiles were quantified in four replicates for every timepoint, and state. TR is in full colors, MO is transparent.

**Figure S2:**
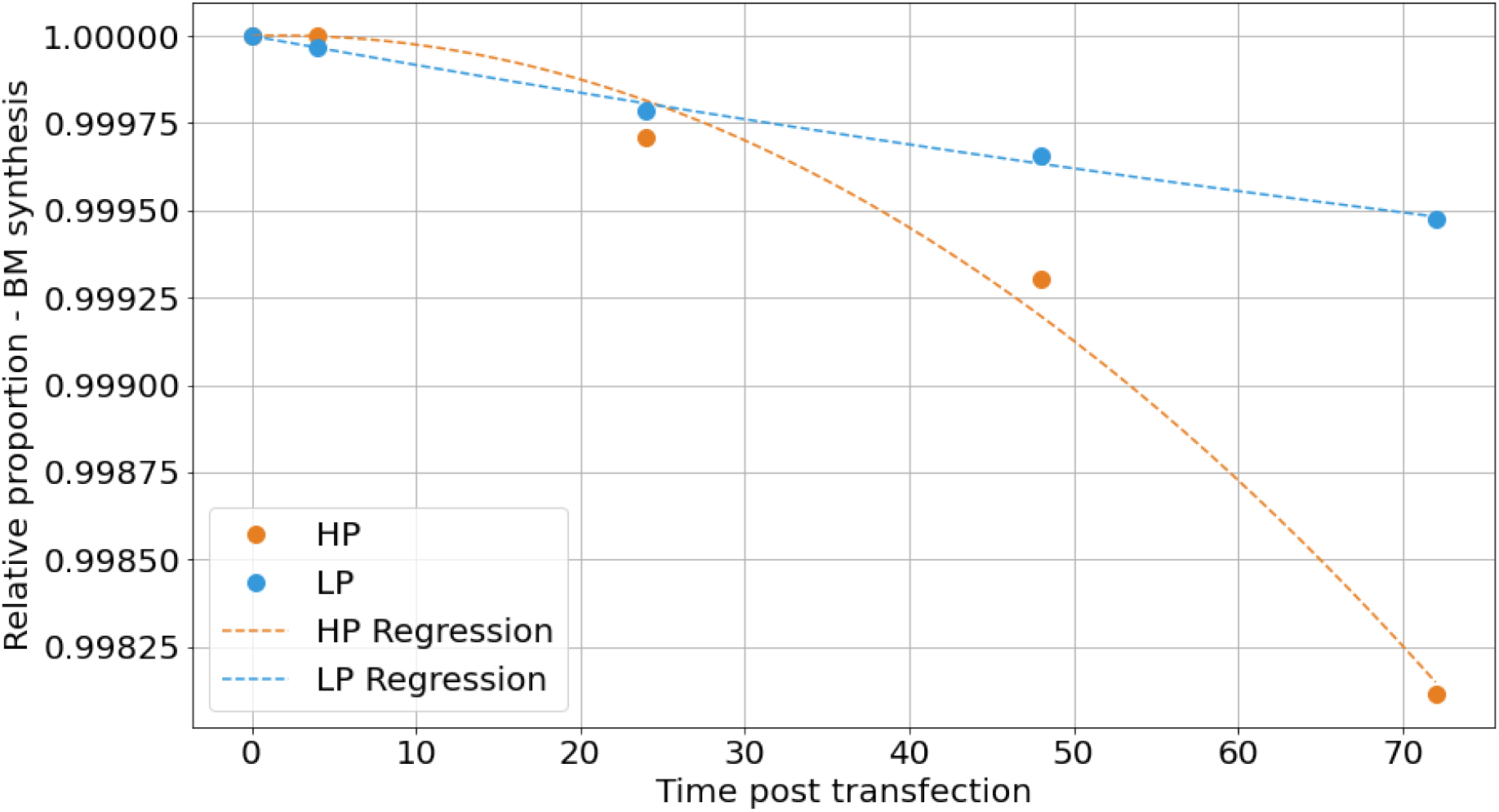
Relative proportion of BM synthesis when considering experimental AAV production. The HP strain (orange) showed a higher proportion of resources used for AAV production, especially from 48 HPT, compared to the LP strain. However, in both strains over 99.8% of the resources were used for growth and not for production.

**Figure S3:**
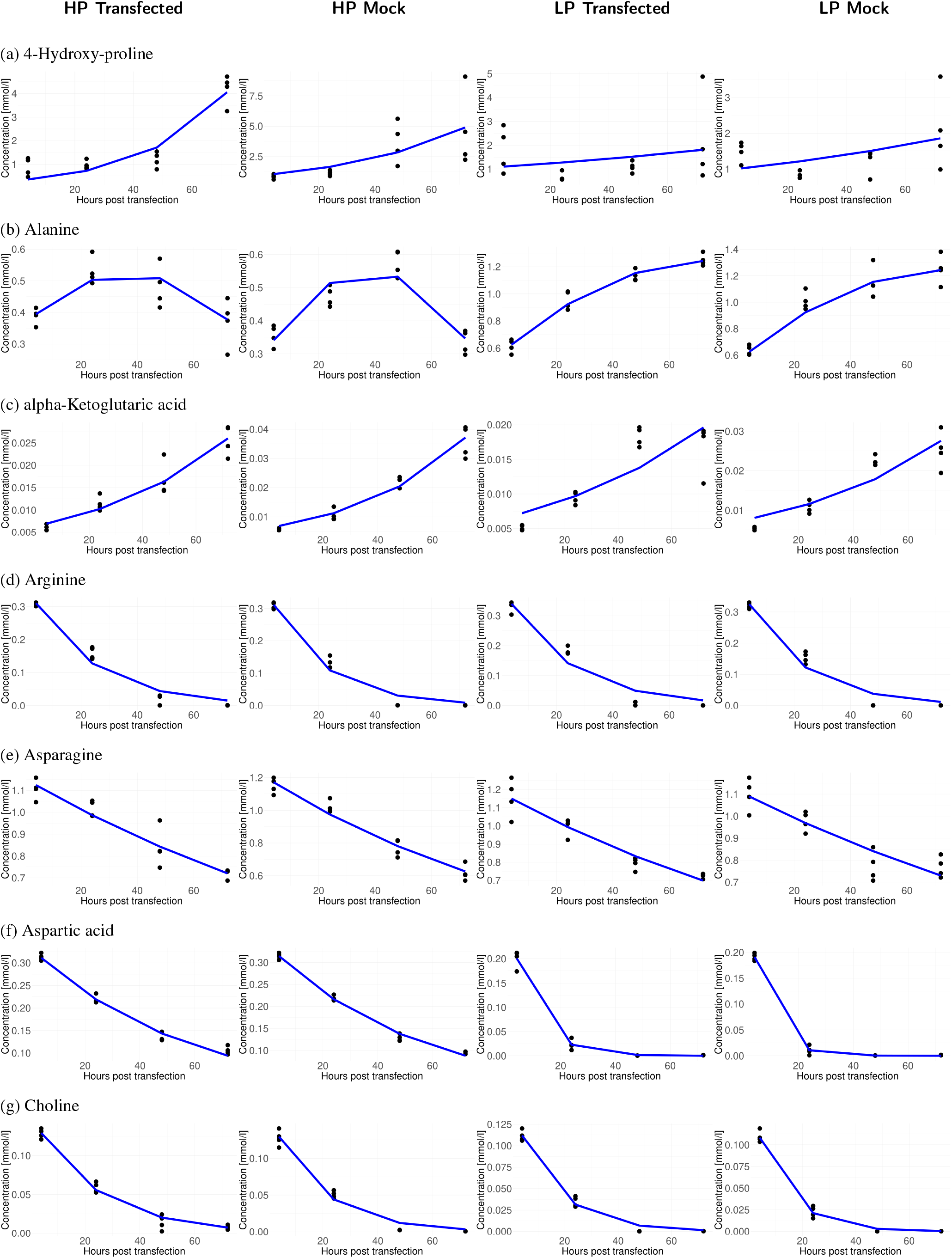
Metabolic profiles from spent media analysis for metabolites including 4, 24, 48 and 72 HPT for HP and LP when producing AAV 8 (transfected) or not (mock). Dots indicate biological repliactes, curves were fitted using the nlrob function in R. See methods for details.

**Figure S4:**
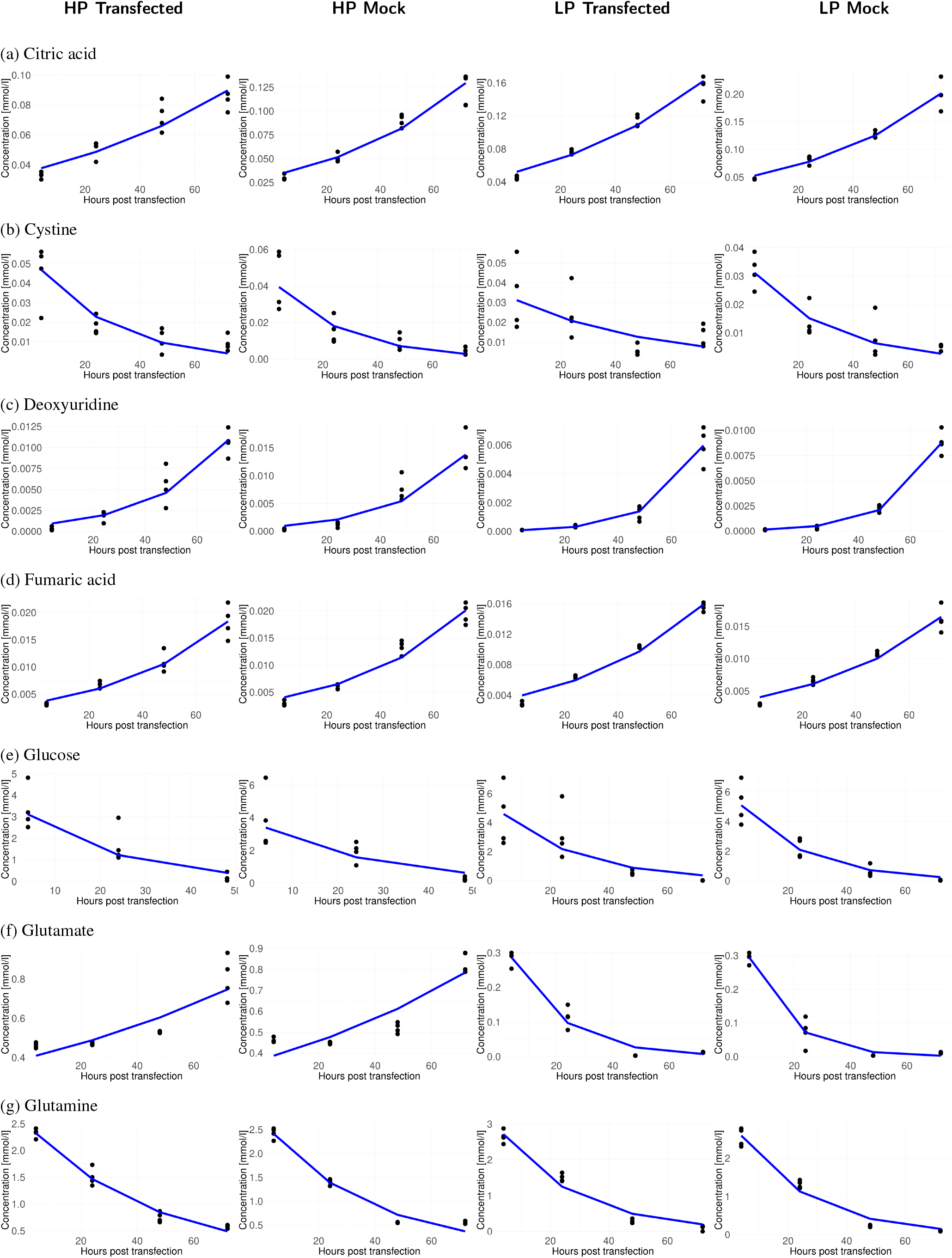
Metabolic profiles from spent media analysis for metabolites including 4, 24, 48 and 72 HPT for HP and LP when producing AAV 8 (transfected) or not (mock). Dots indicate biological repliactes, curves were fitted using the nlrob function in R. See methods for details.

**Figure S5:**
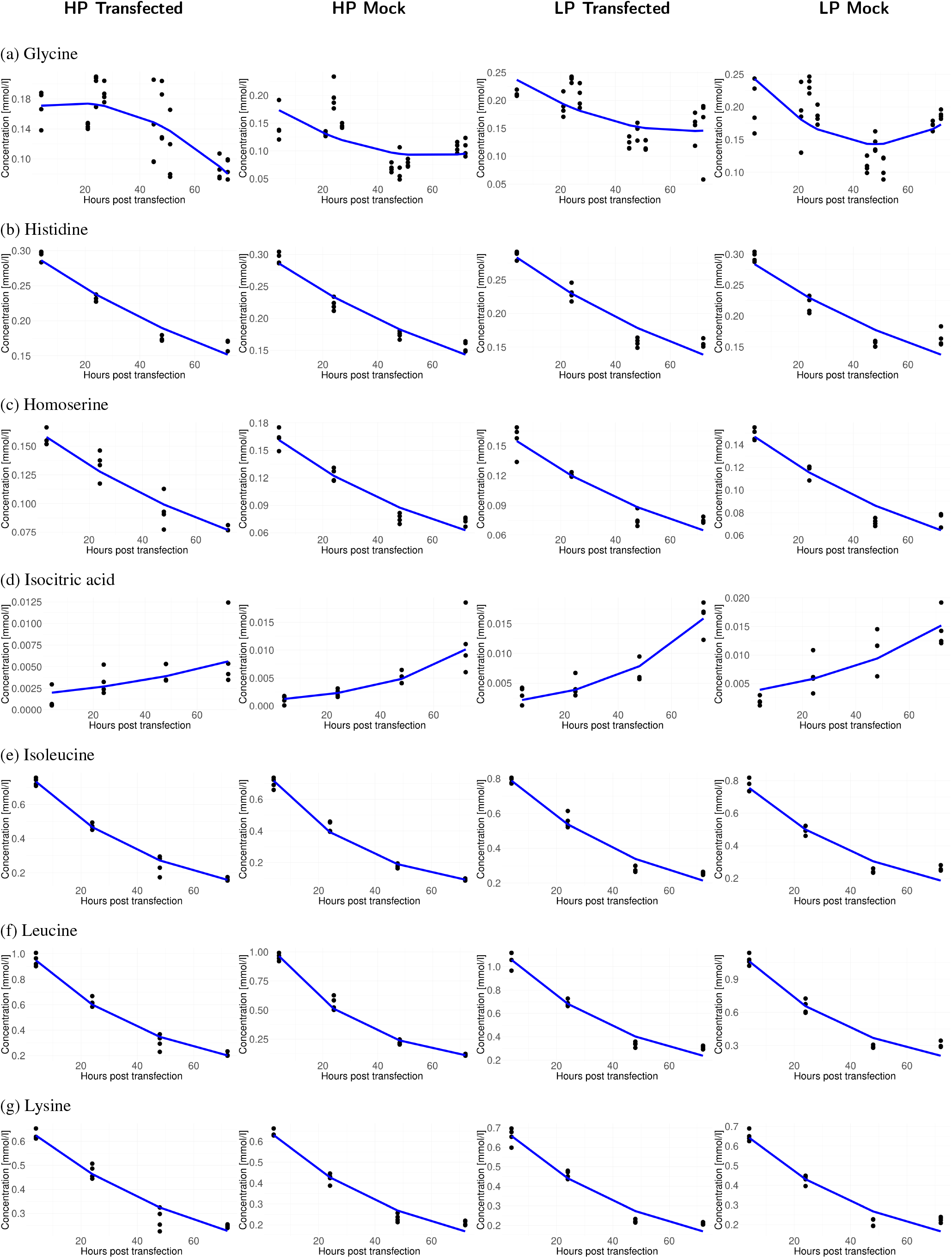
Metabolic profiles from spent media analysis for metabolites including 4, 24, 48 and 72 HPT for HP and LP when producing AAV 8 (transfected) or not (mock). Dots indicate biological repliactes, curves were fitted using the nlrob function in R. See methods for details.

**Figure S6:**
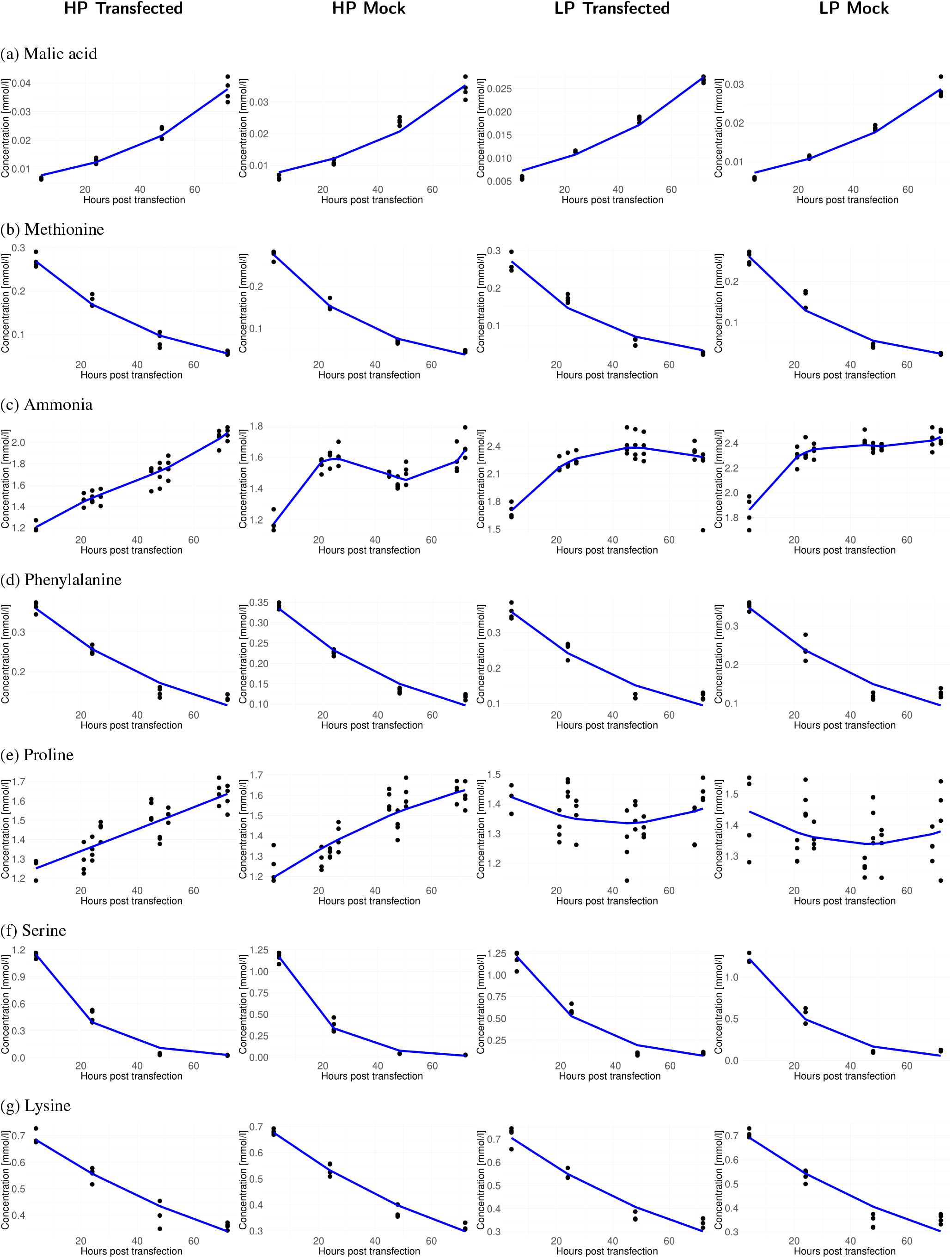
Metabolic profiles from spent media analysis for metabolites including 4, 24, 48 and 72 HPT for HP and LP when producing AAV 8 (transfected) or not (mock). Dots indicate biological repliactes, curves were fitted using the nlrob function in R. See methods for details.

**Figure S7:**
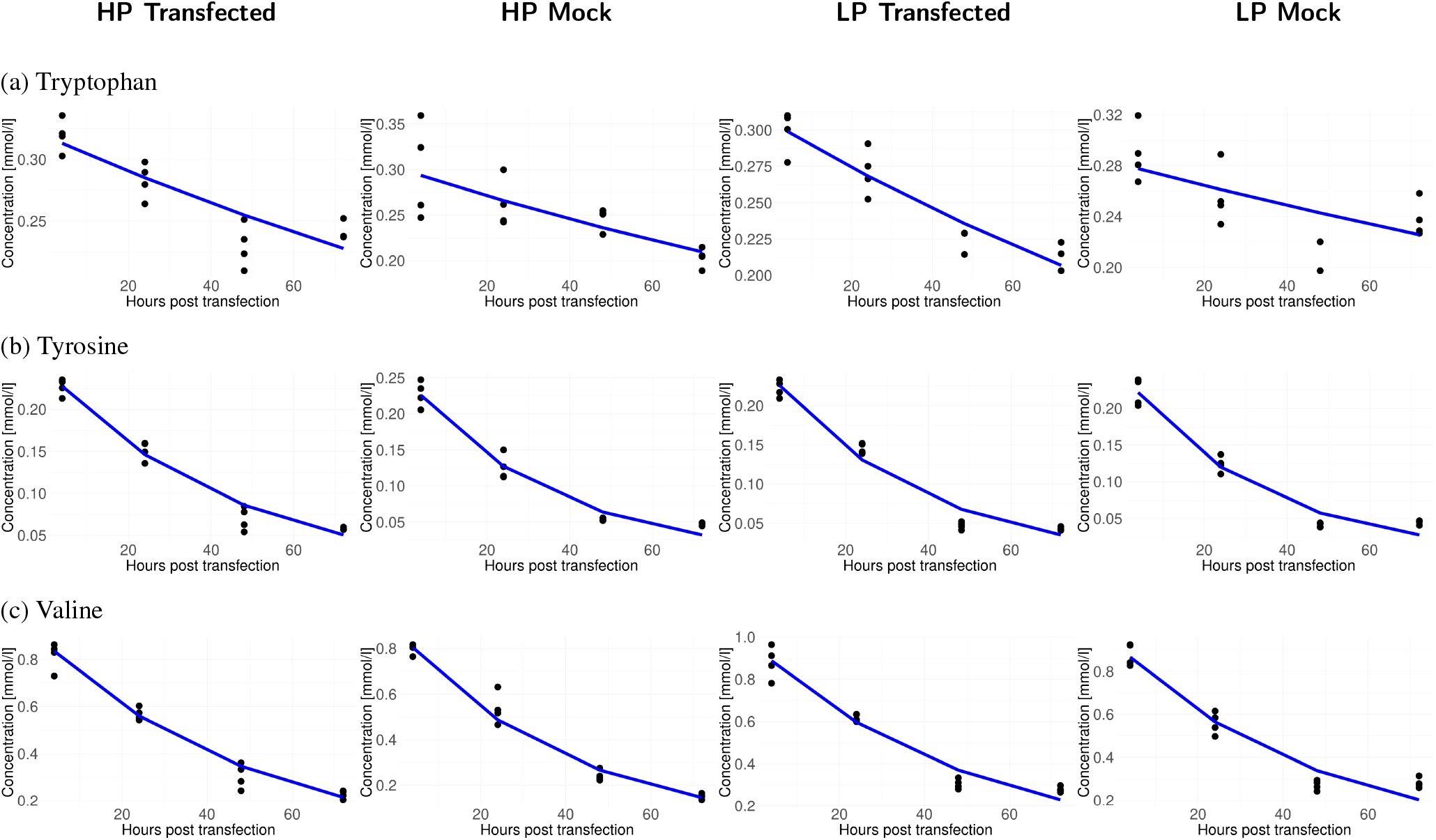
Metabolic profiles from spent media analysis for metabolites including 4, 24, 48 and 72 HPT for HP and LP when producing AAV 8 (transfected) or not (mock). Dots indicate biological repliactes, curves were fitted using the nlrob function in R. See methods for details.

## AAV production reactions

51 ala + 31 arg + 58 asn + 37 asp + 5 cys + 48 gln + 34 glu + 69 gly + 14 his + 24 ile + 53 leu + 32 lys + 11 met + 32 phe + 57 pro + 49 ser + 55 thr + 15 trp + 33 tyr + 30 val + 2215 H_2_O + 2214 GTP → 2214 GDP + 2214 Pi + 1660 H^+^ + 1 VP_1_ 34 ala + 25 arg + 51 asn + 25 asp + 5 cys + 39 gln + 23 glu + 54 gly + 12 his + 23 ile + 37 leu + 22 lys + 10 met + 28 phe + 48 pro + 47 ser + 54 thr + 12 trp + 27 tyr + 25 val + 1804 H_2_O + 1803 GTP → 1803 GDP + 1803 Pi + 1352 H^+^ + 1 VP_2_

30 ala + 21 arg + 49 asn + 22 asp + 5 cys + 34 gln + 20 glu + 45 gly + 12 his + 22 ile + 35 leu + 17 lys + 10 met + 27 phe + 35 pro + 40 ser + 50 thr + 12 trp + 27 tyr + 22 val + 1605 H_2_O + 1605 GTP → 1605 GDP + 1605 Pi + 1204 H^+^ + 1 VP_3_ 5 VP_1_ + 5 VP_2_ + 50 VP_3_ → 1 capsid

1175 ATP + 1175 GTP + 1175 CTP + 1175 TTP → transgene

1 transgene + 1 capsid → 1 full capsid (nucleus)

1 full capsid (nucleus) → 1 full capsid (cytoplasm)

1 full capsid (cytoplasm) → 1 full capsid (extracellular space)

1 full capsid (extracellular space) →

